# Shared patterns of population genomic variation and phenotypic response across rapid range expansions in two invasive lady beetle species

**DOI:** 10.1101/2023.01.13.523993

**Authors:** Angela G. Jones, John. J. Obrycki, Arun Sethuraman, David W. Weisrock

## Abstract

Non-native lady beetle species have often been introduced, with variable success, into North America for biological control of aphids, scales, whiteflies, and other agricultural pests. Two predatory lady beetle species, *Propylea quatuordecimpunctata* and *Hippodamia variegata*, both originating from Eurasia, were first discovered near Montreal, Quebec, in North America in 1968 and 1984, respectively, and have since expanded into northeastern North America and the midwestern United States. In this study, we estimate the range-wide population structure, establishment and range-expansion, and recent evolutionary history of these species of non-native lady beetles using reduced representation genotyping-by-sequencing via ddRADseq. In addition, we quantified the responses to a key abiotic factor, photoperiod, that regulates adult reproductive diapause in these two species and may influence their latitudinal distribution and spread in North America. Our analyses detect (1) non-significant genetic differentiation and divergence among North American populations, (2) evidence of reduced contemporary gene flow within the continental US, (3) significant phenotypic differences in diapause induction despite genetic similarities across sampled populations.

## Introduction

Understanding how species establish new populations in previously unoccupied geographic regions is important in predicting future evolutionary success, especially during times of increased environmental change. This could apply to species that are undergoing rapid shifts in their geographic ranges, often as a response to a changing environment and global climate change (e.g., Collevatti, Nabout, & Diniz Filho, 2011; Karban & Strauss, 2004; Melles, Fortin, Lindsay, & Badzinski, 2011), and also to species that have been intentionally introduced to novel geographic ranges (e.g., Gillis & Walsh, 2017; Ochocki & Miller, 2017; Osborne, Diver, & Turner, 2013). For this latter category, the general factors responsible for, or resulting from, the establishment and expansion of exotic species (e.g., genetic diversity, population structure, local adaptation) are often difficult to discern (Lodge, 1993). For example, some introduced species display significant population structure within twenty years of establishment (LaRue, Ruetz, Stacey, & Thum, 2011; Wang et al., 2017), while others have remained genetically undifferentiated across their invasive range (Tsutsui, Suarez, Holway, & Case, 2000).

The evolutionary potential for nascent populations to survive in new locations is expected to be influenced by many factors, including environmental conditions in the non-native ranges, and the plastic or adaptive phenotypic responses to novel environments (Jardeleza, Koch, Pearse, Ghalambor, & Hufbauer, 2022). Often, one of the first steps in understanding how populations expand into new ranges involves studying the spatial distribution of genetic variation as a means to understand how evolutionary processes structure populations and their phenotypic variation. For example, multiple introductions, resulting in increased genetic diversity and higher adaptive potential have been described in several species (e.g., brown anoles, Kolbe, Larson, & Losos, 2007; American minks, Zalewski et al., 2011; thiarid snails, Facon, Pointier, Jarne, Sarda, & David, 2008; seven-spotted lady beetles, Kajita, O’Neill, Zheng, Obrycki, & Weisrock, 2012; summarized in Dlugosch & Parker, 2008). Similarly, numerous studies have described plasticity in invasive species that have shifted their ranges or microgeographic niches because of changing environments in their previously native ranges (e.g., *Oxalis* geophytes, González-Moreno, Diez, Richardson, & Vilà, 2015; cane toads, Alex Perkins, Phillips, Baskett, & Hastings, 2013; wood frogs, Arietta & Skelly, 2021; summarized in Moran & Alexander, 2014). With increased access to large-scale genomic data and complementary phenotypic and geographical data, studying invasive species has turned an important corner in elucidating: (1) genomic variants associated with adaptations (Stuart et al., 2021), (2) establishing patterns of founder effects and drift across the genome to inform gene drives in controlling invasive species (Oh et al., 2021), and (3) understanding the adaptive potential to previously uninvaded locations using genomic and species distribution models (Mathieu-Bégné et al., 2021).

Biological control organisms such as predatory lady beetles (Coleoptera: Coccinellidae) provide an ideal study system to investigate the establishment, success, and future adaptive potential of invasive species (Sethuraman, Janzen, Weisrock, & Obrycki, 2020; Szűcs, Vercken, Bitume, & Hufbauer, 2019). Numerous species of lady beetles have been introduced to North America for biological control of agricultural pests, with several becoming established (Gordon, 1985; Goryacheva & Blekhman, 2017; Obrycki & Kring, 1998; Sethuraman, Janzen, Rubio, Vasquez, & Obrycki, 2018; Sethuraman et al., 2020) and invasive in their introduced ranges.

For example, based on an analysis of genetic variation in mitochondrial DNA (COI), Kajita et al. (2012) concluded that the distribution of the non-native lady beetle, *Coccinella septempunctata*, in the USA resulted from multiple human introductions and natural range expansion from established populations in the eastern USA. Similar studies in *Harmonia axyridis* (Lombaert et al., 2010, 2011) indicate patterns of successful global invasion from their native range in Eurasia, with numerous range expansions and admixture in Europe, particularly traced to a single source population in Eastern China. On the flipside, releases of hundreds of thousands of mass-reared individuals of several species of predatory lady beetles for biological control programs have also failed to establish successful populations (Ellis, Prokrym, & Adams, 1999; Prokrym, Pike, & Nelson, 1998).

To date, studies exploring the evolutionary and population genetics of invasive lady beetles have been restricted to allozyme, nuclear microsatellite, or mitochondrial DNA data. More recently, large-scale genotyping by sequencing studies, complemented with relevant phenotypic information, are now yielding important insights into range expansions and establishment in biological control organisms (e.g., Hopper et al., 2019; Stahlke et al., 2021).

Here, we combine these approaches to study the establishment and range expansion of two exotic species of Palearctic lady beetles, *Propylea quatuordecimpunctata and Hippodamia variegata*, for biological control of aphid pests in North America. During the 20^th^ century, agriculturalists attempted multiple times to introduce and establish these two predatory species for biological control of aphid pests (Gordon, 1985, 1987; Prokrym et al., 1998; Rogers, Jackson, Angalet, & Eikenbary, 1972). Collections of these two predatory species were made from a wide range of locations in the Palearctic region, mass-reared and released in several locations in the USA; however, no documented establishment of either species as a result of these efforts has been reported (Prokrym et al., 1998; Rogers et al., 1972). Instead, the establishment of North American populations of both species presumably resulted from passive introduction via shipping in the Saint Lawrence Seaway, with the first known collection of non-introduced individuals near Montreal, Quebec (Dysart, 1988; Gordon, 1987). These species have since spread across northeastern North America, with some populations expanding into the midwestern US (Fig 1, Table S1). However, little is known about: (1) the standing genomic diversity of invasive populations, (2) the potential influence on population structure of the intentional and unintentional releases of both species, (3) phenotypic adaptations to novel environments in both species.

**Figure 1.**
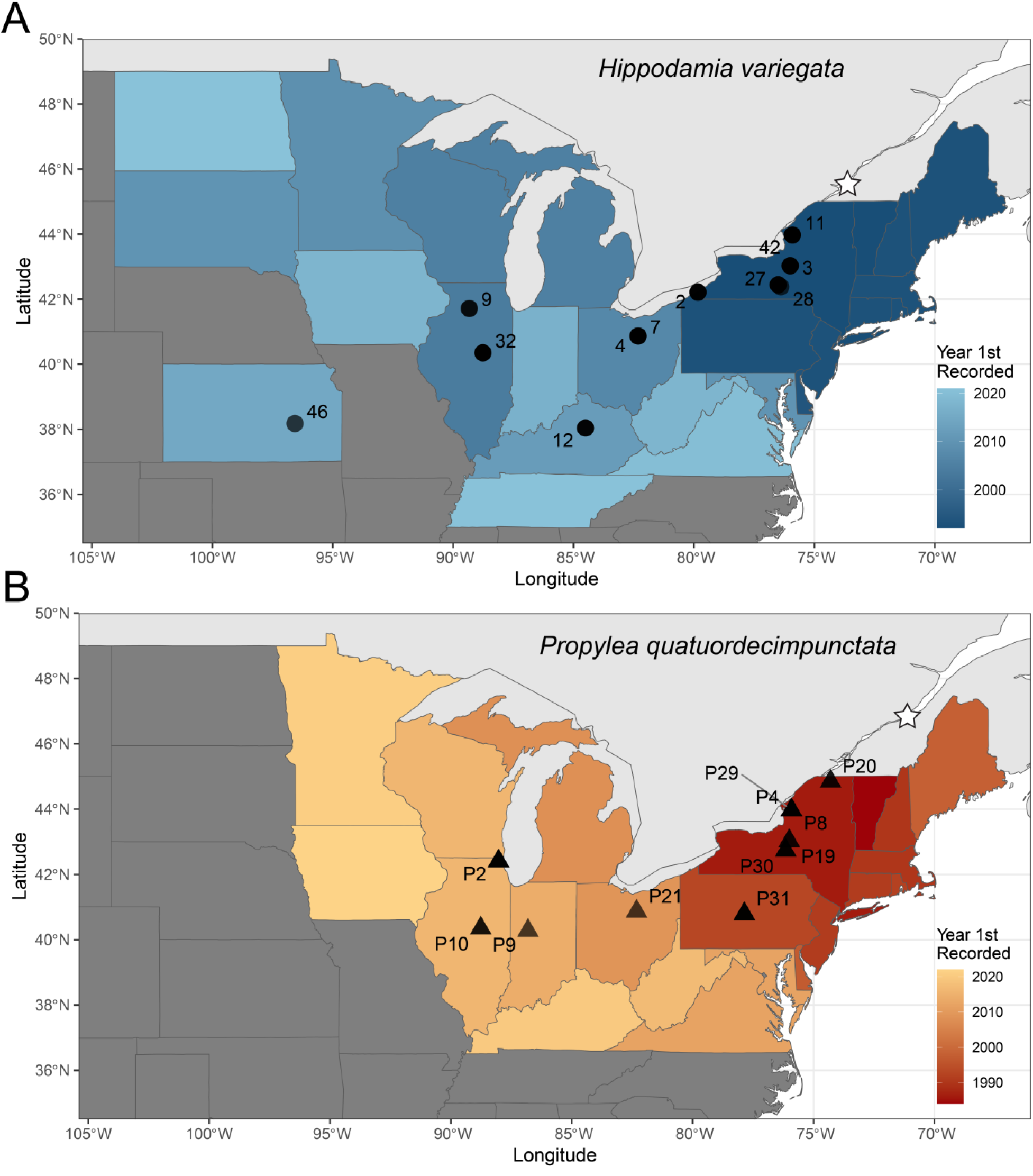
Sampling of (**A)** *H. variegata* and (**B)** *P. quatuordecimpunctata* across their invasive United States ranges. Color gradient denotes year of first recorded specimen in each state. Star icon shows approximate location of North American introduction.

Using a genome-wide dataset of single nucleotide polymorphisms (SNPs) we estimated the population structure of *P. quatuordecimpunctata* (hereafter, P14) and *H. variegata* in their invasive North American ranges through non-spatial and spatially-informed analyses. To complement these genomic studies, we also quantified the responses to photoperiod regulating adult reproductive diapause in both species, aggregating data from *H. variegata* from Obrycki (2018) and a Canadian population of P14 from Obrycki et al. (1993). Previous studies have characterized the responses of North American populations of P14 and *H*. *variegata* to photoperiods as long-day species, in which females under long daylengths avert diapause and short daylengths induce diapause (Obrycki, 2018; Obrycki, Orr, Orr, Wallendorf, & Flanders, 1993). In this study, we compare responses to photoperiods for populations of both species from the same location in North America to determine levels of intra- and inter-specific variation. Considerable phenotypic variation in responses to photoperiod has likely contributed to the invasive of several continents by two non-native lady beetles, *Coccinella septempunctata* and *Harmonia axyridis* (Belyakova, Ovchinnikov, Bezman-Moseyko, & Reznik, 2021; Hodek, 2012; Reznik, Dolgovskaya, Ovchinnikov, & Belyakova, 2015). The response to photoperiods may play a role in the phenology of these species and influence the boundaries of their latitudinal ranges in North America. Specifically, time to onset of diapause is an adaptive plastic trait that influences generation times, fecundity, and overall reproductive fitness of the species (Phoofolo & Obrycki, 2000). We predicted that a plastic response to induction of diapause with range expansion would indicate selection on population fitness.

## Methods

### Sampling, ddRAD Sequencing, and SNP genotyping

We generated SNP data from 37 *H. variegata* individuals from across their known US range (Fig. S1A; Table S2). Also included were four *H. variegata* samples from the Czech Republic, which is within the native range of this species. Our *H. variegata* sampling also included four samples from a population in Australia with unknown origin of establishment and two samples from a population in Chile that likely originated through a biological control program. We generated SNP data from 45 P14 individuals from across their known US range and included four P14 samples from the Czech Republic which is also in the native range of this species (Fig. S1B; Table S3).

We performed double digest restriction-site associated DNA (ddRAD) sequencing using the protocol outlined in Peterson, et al. (2012). Briefly, we extracted DNA from whole individuals using a Qiagen DNEasy tissue kit (Qiagen, Valencia, CA, USA). DNA was quantified with a Qubit 2.0 fluorometer (Thermo-Fisher) and quality-checked with agarose gel electrophoresis. We then used the restriction enzymes *Eco*RI and *Nla*III to digest approximately 250 ng of DNA per individual. Custom inline barcodes and P1/P2 adapters were ligated onto the digested DNA fragments. The uniquely barcoded samples were pooled and size-selected between 473 and 579 bp using a Pippin Prep (Sage Science) machine. Size-selected DNA was then amplified using High-Fidelity DNA Polymerase (Bio-Rad), unique Illumina index sequences, and Illumina sequencing primers for 10 cycles, ensuring that each individual contained a unique set of inline and index sequences. Samples were then pooled into two libraries and sequenced on a single lane of an Illumina HiSeq X at Novogene using 150 bp paired-end (PE) reads. The combined libraries were sequenced using an additional 1% spike of PhiX control library.

We used standard Illumina chastity filtering of our sequence data and demultiplexed reads with STACKS v. 2.53 (Catchen, Hohenlohe, Bassham, Amores, & Cresko, 2013). We then performed downstream processing using ipyrad v. 0.9.19 (Eaton & Overcast, 2020). Reads were then *denovo* assembled using standard settings (described in the code repository). Samples with more than 50% missing data were removed. We filtered at the population level in ipyrad to retain SNPs that were present in at least half of the total populations and in at least half of the total number of individuals in that population. Subsets of U.S. samples were created using the -- branch option. We used VCFTools v. 0.1.16 (Danecek et al., 2011) to sample a single SNP per locus (--thin 5000) resulting in a data set of putatively unlinked SNPs (O’Leary, Puritz, Willis, Hollenbeck, & Portnoy, 2018). To avoid biases in estimation of population structure, we filtered the data in PLINK v. 1.9 (Chang et al., 2015) to remove non-biallelic SNPs as well as SNPs with a minor allele frequency (MAF) less than 0.05 (Linck & Battey, 2019).

### Non-spatially informed population structure

Pairwise F_ST_ between sampling localities was calculated according to Weir & Cockerham (1984) in the R package StaMMP v.1.6.3 (Pembleton, Cogan, & Forster, 2013). Bootstrap replicates (10,000) were used to create a 95% confidence interval for calculated F_ST_ values and test for significance. As a non-model-based test of isolation by distance (IBD) in US populations of *H. variegata* and P14, we used a Mantel test performed in the R package vegan v.1.4.1 with a normalized FST matrix and a Haversine distance geographic matrix (Legendre & Legendre, 2012).

We used two distinct analyses to estimate range-wide subpopulation assignment and admixture of *H. variegata* and P14 individuals. First, we used discriminant analysis of principal components (DAPC) to infer genetic clusters using adegenet (Jombart & Ahmed, 2011; T Jombart, Devillard, & Balloux, 2010). DAPC constructs genetic clusters from a SNP data matrix that have the largest between-group variance and the smallest within-group variance. We used a range of proposed cluster numbers (*K* = 1-10) and used the Bayesian Informative Criterion (BIC) method to estimate the best fit of *K* to the data (Jombart et al., 2010).

Second, we used the program ADMIXTURE v1.3.0 (Alexander & Lange, 2011) to assign individuals to one of *K* genetic clusters. ADMIXTURE jointly estimates subpopulation allele frequencies and admixture proportions under a maximum likelihood framework to quantify deviations from Hardy-Weinberg Equilibrium. We used a range of cluster numbers (*K* = 1-10) across 10 individual replicates and inferred the optimal *K* using a cross-validation (CV) approach (Alexander & Lange, 2011).

### Spatially informed population structure

We used three spatially informed methods to estimate population structure across both species in the US. Spatial principal component analysis (sPCA) was performed in adegenet to estimate genetic dissimilarity on a geographical landscape (Jombart, Devillard, Dufour, & Pontier, 2008). Briefly, sPCA uses Moran’s *I* index to correlate spatial information to genetic allele frequencies. To create the input GENIND file, we used the R tool *vcfR* (Knaus & Grünwald, 2017). Our analyses used a nearest-neighbor connection network accounting for multiple samples at each sampling site (type = 6) with k = 20 and k = 25 neighbors for P14 and *H. variegata*, respectively. This connection network scheme was created to maximize connectivity across spatially distinct samples while accounting for sampling differences between species and potential long-distance dispersal events among sampling sites (Maigret, Cox, & Weisrock, 2020). We estimated the significance of global and local structure with 999 permutations through adegenet (Montano & Jombart, 2017).

Second, we used MEMGENE to identify spatial neighborhoods in genetic distance data. MEMGENE combines Moran’s eigenvalue maps with a regression framework where genetic distances are spatially independent. Samples are mapped based on independent location, and only significant eigenvalues are retained for analysis (Galpern, Peres-Neto, Polfus, & Manseau, 2014).

Lastly, we used conStruct (Bradburd, Coop, & Ralph, 2018) to investigate if genetic structure was due to IBD. ConStruct is a model-based method that simultaneously infers continuous and discrete patterns of population structure by estimating ancestry proportions for each sampled individual from two-dimensional population layer, while estimating the rate at which relatedness decays with geographic distance. This method also allows for cross-validation procedure for model selection, between both spatial and nonspatial models. We used the R package *fields* to create the geographic distance matrix that conStruct utilizes and conducted cross-validation analyses from *K* = 1-10 with 10,000 iterations and 5 replications per *K* value.

### Estimation of migratory history

To estimate evolutionary history of the structured populations identified by our ADMIXTURE analyses across *H. variegata* and P14, we used the Bayesian MCMC tool, BayesAss3-SNPs (Mussmann, Douglas, Chafin, & Douglas, 2019; Wilson & Rannala, 2003). Briefly, BayesAss estimates contemporary migration rates from multilocus genotype data using a Bayesian MCMC method. We performed 48-hour runs of each MCMC, discarding 10% of all sampled states as burn-in, resulting in a total of 4.94e7 states for the *H. variegata* data, and 1.18e7 states for the P14 data. Convergence of the MCMC was assessed using Tracer v1.7.1, and 95% confidence intervals around migration rate estimates were constructed using the marginal posterior density distributions thus obtained.

### Effect of Photoperiod on Preimaginal Development and Diapause Induction

Data for the developmental and reproductive responses to different photoperiod conditions for *H. variegata* were taken from Obrycki (2018). Adult individuals used in that study were sampled from Jefferson County, New York, USA.

New data for P14 were generated using adults collected from Jefferson County, New York, USA (43.98°N, 75.91°W). Individual females or mating pairs were placed in 0.24 L (8 oz.) paper containers (Choice®, http://webstaurantstore.com), maintained at a photoperiod of L:D 16:8 (light:dark), a temperature of 22 ± 1°C, and provided water, a Wheast (GreenMethods.com)-honey mixture, and a daily supply of pea aphids, *Acyrthosiphon pisum* (Harris) (Hemiptera: Aphididae). Eggs were collected daily from 4 to 6 females and placed in L:D 16:8, 14:10, 12:12,10:14, at 22°C ± 1.0°C. F1 offspring were individually reared in glass vials at each photoperiod on *A*. *pisum* and *Ephestia kuehniella* (Zeller) (Lepidoptera: Pyralidae) eggs (Beneficial Insectary, Redding, CA).

Pairs of F1 adults were placed in 0.24 L paper containers at the same larval photoperiod, provided water, a Wheast-honey mixture, and pea aphids. Pairs were fed and maintained at each photoperiod for 120 days, when the experiment was ended. The date of first oviposition was recorded for each female. If a female died, the date of her death was recorded. If a male died, a male from the same L:D condition was used as a replacement. Voucher specimens of P14 are accessioned in the University of Kentucky Insect Museum.

The length of the pre-oviposition period (days) was recorded to quantify the proportion of P14 females at a given photoperiod that was in diapause and estimate the duration of diapause in females at the four photoperiods. A prolonged pre-oviposition period was observed in some females at long daylengths (L:D 16:8) and excess prey availability, conditions that typically do not induce diapause. The diapause or non-diapause condition of each P14 female at the four photoperiods was based on twice the median pre-oviposition period (days) observed at L:D 16:8. This type of classification has been used to separate females into diapause and non-diapause groups in previous studies of adult reproductive diapause in predatory Hemipterans (Ruberson, Shen, & Kring, 2000; Ruberson, Yeargan, & Newton, 2001) and coccinellids (Obrycki, 2018, 2020, 2022).

### Photoperiod Statistical Analyses

Egg, larval, pupal, and pre-imaginal (egg to adult) developmental times (days) for P14 were compared among the four photoperiods using one-way ANOVA (JMP Pro 14.0.0). The developmental times for P14 and *H. variegata* populations from Jefferson County (*H. variegata* data from Obrycky, 2018) were compared in this study using a two-way ANOVA (JMP Pro 14.0.0). The days from female eclosion to first oviposition (pre-oviposition period) at each photoperiod was compared within each species and between species for populations collected in Jefferson County using event-time analysis (JMP Pro 14.0.0). Females that died or did not oviposit within 120 days, the duration of the experiment, were censored, because the pre-oviposition period for these individuals was not measured. A non-parametric log-rank analysis of the response to photoperiod within each population was used to examine if the pre-oviposition period varied among females at each photoperiod (JMP Pro 14.0.0). Comparisons of responses to the four photoperiods between P14 from Jefferson County to P14 from Montreal, Canada were also conducted using the non-parametric log-rank test (JMP Pro 14.0.0). Data for the Montreal, Canada population are from Obrycki et al. (1993).

## Results

### Sampling, ddRAD Sequencing, and SNP genotyping

We generated a total of ~996 million 150 bp PE reads with a mean of 10,379,052 reads per individual. After demultiplexing and standard Illumina filtering, an average of 82.3% of reads were retained per individual. Two sub-libraries sequenced poorly and retained an average of 25.1% of reads after demultiplexing. The average percentage retained excluding those poorly sequenced was 96.6%. After filtering, we recovered genotypes for 70 individuals (41 *H. variegata*; 29 P14) from 26 localities (16 *H. variegata*; 10 P14). Filtering by a minor allele frequency of 0.05 resulted in a total of 634 loci containing SNPs in *H. variegata* and 2196 loci containing SNPs in P14. Data sets restricted to US samples resulted in a total of 876 loci containing SNPs in 32 *H. variegata* individuals and 2359 loci containing SNPs in 25 P14 individuals.

### Non-spatial population structure

Pairwise F_ST_ analyses of *H. variegata* revealed significant (p<0.05) genetic distance between continental groups (Table S4). In addition, we detected small (F_ST_ < 0.1) but significant effects between some North American populations which roughly follow an east-west cline. Parallel analysis of P14 again revealed significant separation by continent and small differences along an east-west pattern between North American populations (Table S5). Mantel tests did not detect significant signatures of IBD in North American samples of *H. variegata* (*p* = 0.25) or P14 (*p* = 0.39). Neither DAPC nor ADMIXTURE identified distinct geographic genetic clusters in the North American range of either species. The BIC value of DAPC and the cross-validation of ADMIXTURE were lowest for *K* = 1 (Fig. S2). When *K* = 2 was considered for *H. variegata*, ADMIXTURE (Fig. 2A) and DAPC (Fig. S2) assignment plots supported a separation of North American and Czech samples from Australian and Chilean samples. *K* = 2 assignment plots for P14 clustered the Czech samples with an assortment of North American samples (Fig. 3A), but without any clear geographic patterning to the North American samples.

**Figure 2.**
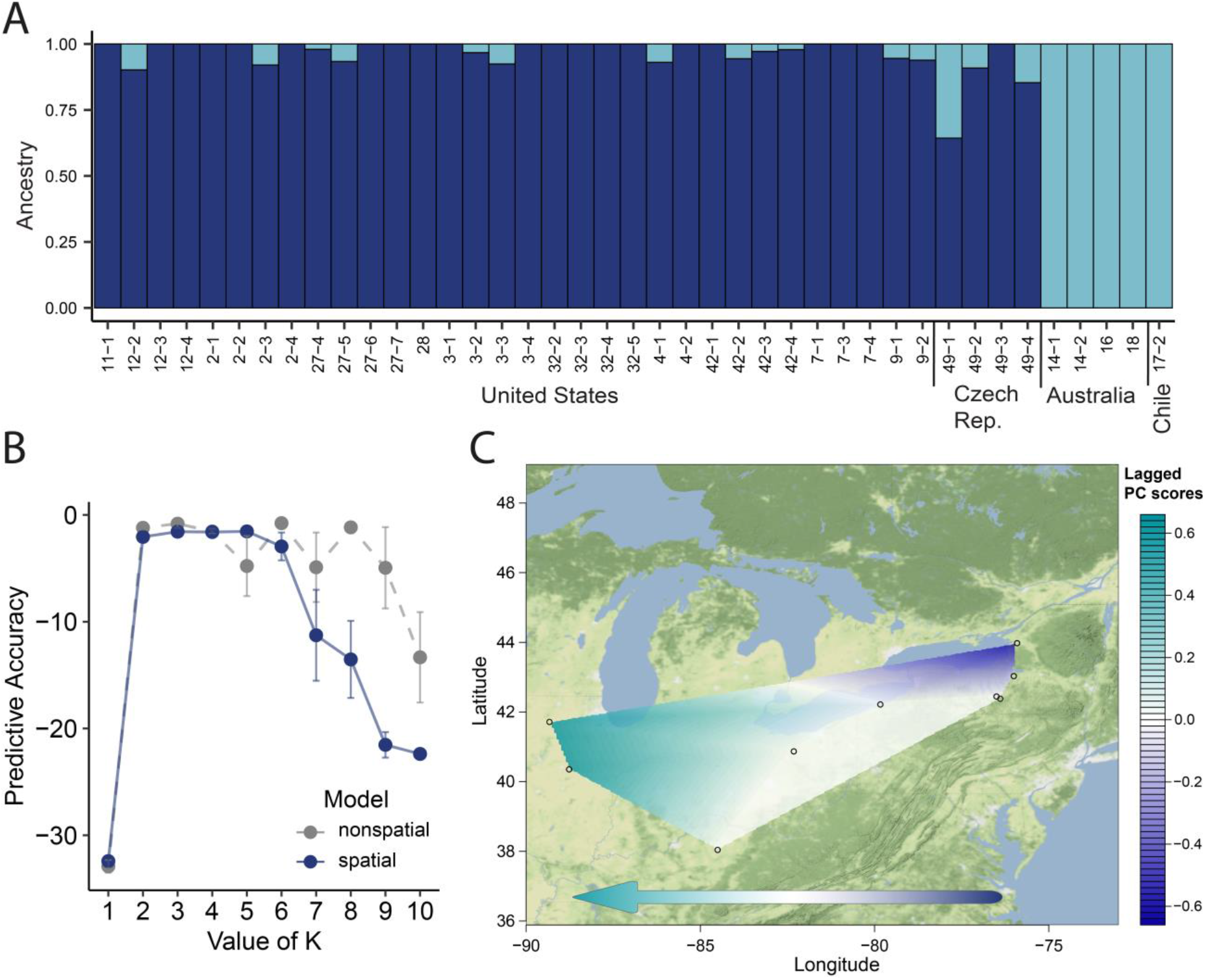
Population structure analyses of *H. variegata* in the United States. **(A)** ADMIXTURE assignment plot for *K*=2. **(B)** conStruct cross-validation results for *K*=1-10 in both spatial and non-spatial analyses. **(C)** sPCA lagged principal component scores across populations.

**Figure 3.**
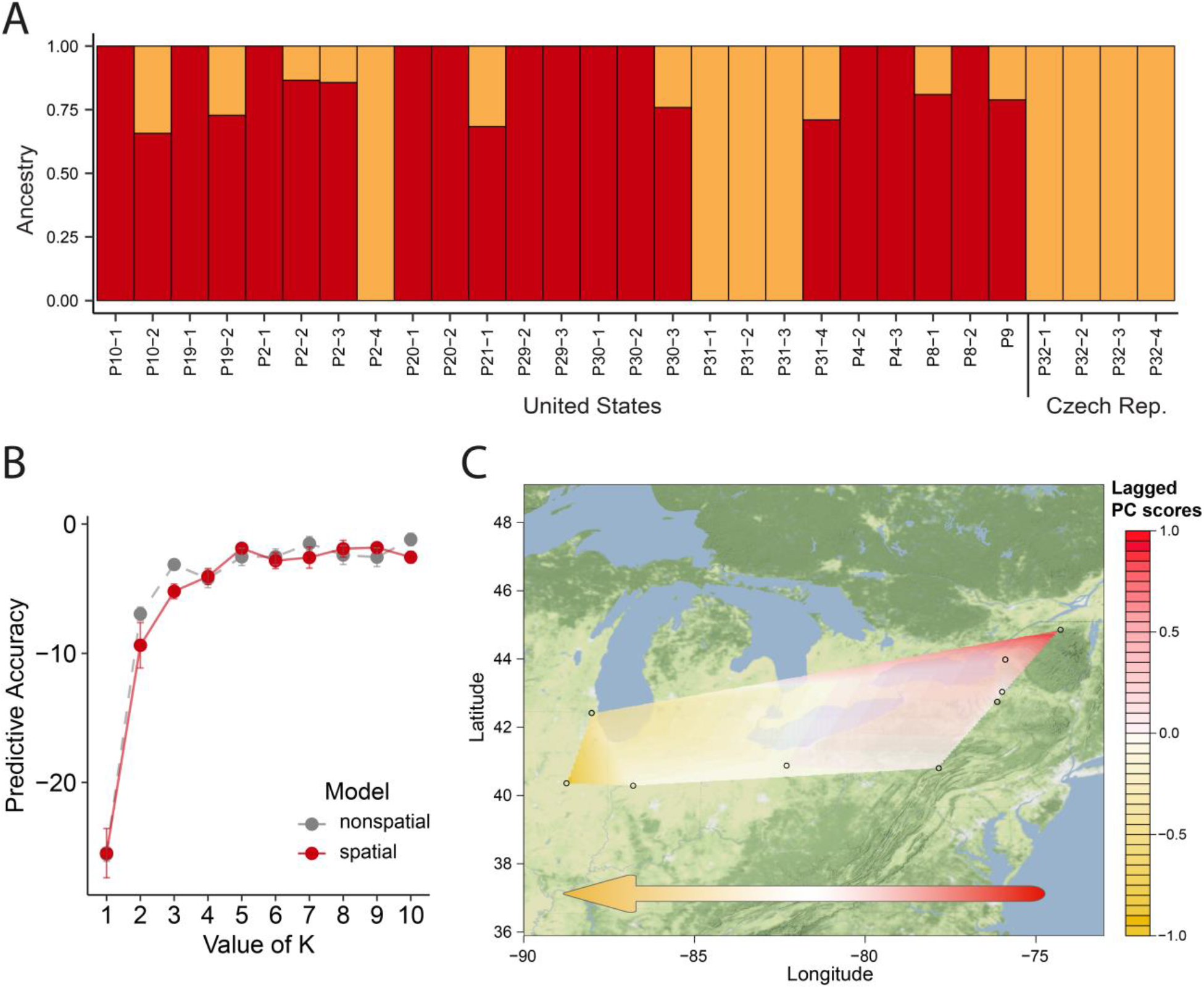
Population structure analyses of *P. quatuordecimpunctata* in the United States. **(A)** ADMIXTURE assignment plot for *K*=2. **(B)** conStruct cross-validation results for *K*=1-10 in both spatial and non-spatial analyses. **(C)** sPCA lagged principal component scores across populations.

### Spatial population structure

MEMGENE did not detect any significant eigenvectors in either *H. variegata* and P14, and thus no results were produced. Cross-validation conStruct analyses of both *H. variegata* (Fig. 2B) and P14 (Fig. 3B) indicated minor differences in the predictive accuracy of spatial and nonspatial models, suggesting that geographic distance does not significantly contribute to genetic differences. Interestingly, conStruct cross-validation analyses indicate that a *K* = 2 has the highest predictive accuracy of tested *K*s in both *H. variegata* and P14. However, further investigations revealed that only a single *K* is notably contributing to spatial model layers (Fig. S3). Thus, these results do not detect a signature of population structure affected by IBD.

For the U.S. dataset of *H. variegata*, sPCA detected an eigenvector that denoted an east-west patterning of genetic differentiation (Fig. 2C). However, neither global (*p* = 0.61) nor local structure (*p* = 0.17) was significant. SPCA of P14 yielded similar results with a single eigenvector delimiting an east-west division (Fig. 3C). Again, there was no significant structure on the global (*p* = 0.96) or local (*p* = 0.37) scales.

### Contemporary Migratory History

Estimates of contemporary migration using BayesAss3-SNPs reaffirmed our admixture analyses using STRUCTURE and ADMIXTURE, with most migration estimated only within sampled populations, and negligible migration rates between populations (Tables S6, S7) in both populations of P14 (*K* = 2 of individuals in the USA, and Czech Republic) and *H. variegata (K* = 3, separated into individuals in the USA, Czech Republic, and Australia + Chile). All MCMC runs were assessed for convergence and mixing using Tracer v.1.7.1 by observing the trace-plots for migration rates (to ensure mixing), large effective sample sizes (ESS), and autocorrelations.

### Photoperiodic effects on preimaginal development and diapause induction

Photoperiod significantly influenced egg, larval, pupal and total pre-imaginal development of P14 collected in Jefferson County, New York (Tables S8, S9). The adult reproductive responses to photoperiod, as measured by the pre-oviposition period varied significantly among the four photoperiods (Table S10, P14 analyses within row). The median pre-oviposition period for P14 females at L:D 16:8 was 10 days; no oviposition was observed at L:D 14:10, 12:12 or 10:14).

### Intraspecific and interspecific comparisons of responses to photoperiod

The responses of two P14 populations from Montreal, Quebec, Canada and Jefferson County, New York, USA to the four constant photoperiods were similar (Table S11: analyses within columns). The non-parametric log-rank analysis of pre-oviposition periods indicated significant differences in responses by P14 and *H. variegata* collected in Jefferson County, New York to L:D 14:10 and L:D 12:12, but similar responses to L:D 16:8 and L:D 10:14 (Table S12, analyses within rows). At each of the four constant photoperiods, a similar percentage of female P14 and *H. variegata* from Jefferson County, NY and P14 (P14CAN) females from Montreal, Quebec, Canada were induced into diapause (Table S12).

## Discussion

Here we analyze genetic diversity, distribution, and patterns of phenotypic variation in two different invasive species of lady beetles utilized in biological control, with shared patterns of importation and augmentation history. Our analyses indicate seemingly rapid Westward expansion of both species, with little discernible population structure evolving across its non-native range. In contrast, experimental analyses of phenotypic variation in photoperiod response in development and diapause induction identified significant differences between populations within both species, suggesting the potential for either local adaptation or phenotypic plasticity.

### Genetic differences in native and introduced populations

Non-spatial estimates of population structure showed little difference between European and North American samples in both species. This result parallels previous allozyme results in *H. variegata* (Krafsur, Obrycki, & Nariboli, 1996) and P14 (Krafsur & Obrycki, 1996), suggesting minimal significant genome-wide genetic differentiation between populations from the native and introduced ranges. In addition, no significant differences have been detected in physiological mechanisms such as photoperiod and net reproductive rates between North American and European samples of P14 (Obrycki et al., 1993). This suggests that all non-native populations of both species in North America were derived from a small number of founders that underwent a bottleneck early in their importation, with little successful augmentation from secondary introductions, as discovered in other invasive lady beetle species (e.g. *Harmonia axyridis* – Lombaert et al., 2014, 2011; *Coccinella septempunctata* – Kajita et al., 2012).

In contrast, Chilean and Australian samples of *H. variegata* appear to be genetically distinct from European and North American samples, suggesting that the founding individuals of Chile and Australia originated from a different source population than those of North America. During the 1970s, *H. variegata* from South Africa were released into Chile; the origins of the Australian populations are not known (Franzmann, 2002; Grez, Viera, & Soares, 2012). Previous studies have suggested eastern Asia as the origin of *H. variegata* due to higher levels of genetic diversity found in that region (Krafsur et al., 1996). Our analyses do not include samples from Asia, but further investigation could determine the source populations of invasions into Chile and Australia.

Within the U.S.A, we uncovered weak signatures of gene flow across the ranges of both *H. variegata* and P14. sPCA eigenvectors support an east-west divide in both species, indicating signatures of isolation by distance (IBD). These differences are likely due to genetic drift associated with increasing geographic distance (Krafsur et al., 1996). Non-spatial analyses and IBD-informed analysis through conStruct however were unable to detect any significant population structure in the invasive ranges of both P14 and *H. variegata*.

While our results do not show significant genetic structure within the invasive range, this is perhaps not entirely unexpected. *H. variegata* and P14 were both introduced to North America within the last 55 years. Within this time, these species have rapidly expanded and have proved successful in colonization of new territory. Invasive species are often able to spread quickly with few genetic signatures of divergence (Eyer et al., 2018; Tsutsui et al., 2000). This is particularly true of coccinellid species where single females establish new colonies when there is sufficient prey density (Krafsur & Obrycki, 1996). These colonizing events cause successive founder effects that may be responsible for the lack of significant structure and within population genetic diversity in the U.S. populations of *H. variegata* and P14.

Our analyses of spatial structure and dispersal indicate that both species are showing natural dispersal from their source populations in a generally western direction. *H*. *variegata* disperses at a more rapid rate than P14; the former species was first reported in Canada in 1984, whereas P14 was first collected in Canada in 1968, but *H*. *variegata* currently has a larger North American distribution. *H*. *variegata* has also recently been found in Oregon, in the western U.S.A (Jessie, Reich, & Mc Donnell, 2020). Additional study across a broader geographical range is required to determine that this western collection of *H*. *variegata* does not represent a new introduction of this species.

### Photoperiod response during a rapid range expansion

While our analyses of genetic structure indicated no significant differences among invasive populations in the USA, significant phenotypic variability in invasion-related traits in P14 and *H. variegata* indicate that the expanding distribution of both species may be related to their seasonal phenology, characterized by their responses to abiotic factors, including photoperiodic induction of diapause, in their new North American habitat. Phenotypic plasticity (Shearer et al., 2016), pre-adaptation (Reznik et al., 2015), and/or post-colonization evolution (Bean, Dalin, & Dudley, 2012) in responses to the abiotic factors temperature and photoperiod have contributed to the geographic expansion of invasive insect species. North American populations of P14 and *H*. *variegata* from the northeastern USA and southeastern Canada generally responded similarly to four constant photoperiods at 22°C, with significant number of ovipositing females, and diapause induction across diurnal periods. However, in our comparisons of P14 populations from Montreal, Quebec, Canada (Obrycki et al., 1993) to the population from Jefferson County, New York, USA in this study, we did not detect any significant (p > 0.05) post-colonization differences in the photoperiodic responses of P14. This pattern of longer-day mediated oviposition is seen in both *H*. *variegata* and *P*. *quatuordecimpunctata*, in which females under long daylengths avert diapause and short daylengths induce diapause. This photoperiodic response has been documented for several predatory lady beetles that diapause as adults (e.g., *Adalia bipunctata, Coccinella septempunctata, Harmonia axyridis, Hippodamia convergens*, and *Hippodamia parenthesis*) (Hodek, 2012; Obrycki, 2018, 2020; Obrycki et al., 1983). Previous studies suggest that reproductive diapause in *H*. *variegata* might end by January and that initiation of vernal activity may be regulated by temperature (Obrycki, 2018). Data for P14 presented in this study indicates that this species may have a prolonged diapause regulated by increasing daylengths in the spring. A possible explanation for the more widespread distribution of *H*. *variegata* is the opportunistic production of two generations/year compared to a single generation of P14 (Day & Tatman, 2006).

Based on our current understanding of the genomic and ecological characteristics of the North American populations of *H. variegata* and P14, it appears that both species were unintentionally introduced into southeastern Canada near Montreal, Quebec in the latter half of the 20^th^ Century. Multiple geographic populations of both species were mass reared and released in large numbers for biological control of aphid pests in the U.S.A., but no established populations were ever confirmed based on these releases. Our genomic analysis indicates little genetic structure in the North American populations of *H. variegata* and P14 supporting the argument of natural spread from the original accidentally established Canadian populations.

### Summary

In summary, our findings indicate shared patterns of introductory history of two species of invasive lady beetles, leading to serial founder effects, and reduced phenological response among non-native populations in the United States. Our stud thus supports a rapid response to shorter diurnal cycles in lower latitudes, either through adaptation or phenotypic plasticity, and a generally westward invasion by both species, despite low genomic diversity and population structure. This suggests that the success of invasive species need not necessarily be mediated by higher genomic diversity of founding populations, or bridgehead effects from secondary augmentations, but perhaps contingent on standing genetic diversity and adaptive evolution to local environments.

## Acknowledgments

This work was supported by NSF ABI 1564659, NSF CAREER 2042516 to AS. This work was funded by the National Institute of Food and Agriculture, U.S. Department of Agriculture, Hatch Program under accession number 1008480 and funds from the University of Kentucky Bobby C. Pass Research Professorship to JJO. This research was supported in part by a Research Support Grant from the University of Kentucky Office of the Vice President for Research to DWW and JJO. This research includes calculations carried out on HPC resources supported in part by the National Science Foundation through major research instrumentation grant number 1625061 and by the US Army Research Laboratory under contract number W911NF-16-2-0189. We thank the following for specimens: A. Slipinski, Australian National Insect Collection, CSIRO, Canberra, Australia; A. Grez, Department of Animal Biological Sciences, Universidad de Chile, Santiago,Chile; Y. Kajita, University of North Carolina, Chapel Hill, NC, USA.

## Data Availability

Sequence data will be available on the NCBI Sequence Read Archive after publication (BioProject accession PRJNA911772). Input files for all population genetic analyses will be available via figshare after publication (10.6084/m9.figshare.21896991). Scripts used for all analyses will be available via figshare after publication (10.6084/m9.figshare.21897075).

## Author Contributions

Designed research: A.G.J., A.S., J.J.O, and D.W.W.

Performed research: A.G.J., A.S., and J.J.O

Analyzed data: A.G.J., A.S., and J.J.O

Wrote the paper: A.G.J., A.S., J.J.O, and D.W.W.

## Notes

https://doi.org/10.6084/m9.figshare.21899385

### Competing Interest Statement

The authors have declared no competing interest.

