## Supplementary Material for "Shared patterns of population genomic variation and phenotypic response across rapid range expansions in two invasive lady beetle species"

### Table of Contents:


| | |
|---|---|
| **Table S1** | Page 2 |
| **Table S2** | Page 3 |
| **Table S3** | Page 4 |
| **Table S4** | Page 5 |
| **Table S5** | Page 6 |
| **Table S6** | Page 7 |
| **Table S7** | Page 8 |
| **Table S8** | Page 9 |
| **Table S9** | Page 10 |
| **Table S10** | Page 11 |
| **Table S11** | Page 12 |
| **Table S12** | Page 13 |
| **Figure S1** | Page 14 |
| **Figure S2** | Page 15 |
| **Figure S3** | Page 16 |
| **Figure S4** | Page 17 |


**Table S1.** Year and corresponding reference of primary recorded occurrences of *H. variegata* (HV) and *P. quatuordecimpunctata* (P14) in the United States.

| State | HV year | HV reference | P14 year | P14 reference |
|---|---|---|---|---|
| CT | 1993 | Wheeler, et al., 1993 | 1991 | Day, et al., 1994 |
| DE | 1994 | Ellis, et al., 1999 | 1997 | Ellis, et al., 1999 |
| IL | 2004 | Hesler, et al., 2017 | 2014 | Primary data |
| IN | 2017 | GBIF, 2022 | 2012 | Primary data |
| IA | 2018 | Hesler, et al., 2018 | 2022 | GBIF, 2022 |
| KS | 2015 | Primary data | -- | -- |
| KY | 2012 | Primary data | 2020 | GBIF, 2022 |
| ME | 1993 | Ellis, et al., 1999 | 1988 | Day, et al., 1994 |
| MD | 2010 | Losey, et al., 2012 | 2012 | Losey, et al., 2012 |
| MA | 1993 | Wheeler, et al., 1993 | 1990 | Day, et al., 1994 |
| MI | 2005 | Gardiner, et al., 2005 | 2008 | Losey, et al., 2012 |
| MN | 2009 | Heidel, et al., 2011 | 2021 | GBIF, 2022 |
| NH | 1993 | Wheeler, et al., 1993 | 1990 | Day, et al., 1994 |
| NJ | 1993 | Wheeler, et al., 1993 | 1991 | Day, et al., 1994 |
| NY | 1992 | Wheeler, et al., 1993 | 1986 | Day, et al., 1994 |
| ND | 2021 | GBIF, 2022 | -- | -- |
| OH | 2007 | Pavuk, et al., 2007 | 2009 | GBIF, 2022 |
| PA | 1993 | Wheeler, et al., 1993 | 1993 | Day, et al., 1994 |
| RI | 1993 | Wheeler, et al., 1993 | 1992 | Day, et al., 1994 |
| SD | 2010 | Hesler, et al., 2011 | -- | -- |
| TN | 2021 | GBIF, 2022 | -- | -- |
| VT | 1992 | Wheeler, et al., 1993 | 1984 | Day, et al., 1994 |
| VA | 2020 | GBIF, 2022 | 2012 | GBIF, 2022 |
| WV | 2018 | GBIF, 2022 | 2017 | GBIF, 2022 |
| WI | 2005 | Williams, et al., 2009 | 2016 | GBIF, 2022 |

**Table S2.** Sampling localities, GPS Coordinates, and collection information for *H. variegata* individuals.

| Locality Number | Location | Lat | Long | Date Collected | # of Samples Sequenced | # of Samples Dropped |
|---|---|---|---|---|---|---|

| | | | | | | |
|---|---|---|---|---|---|---|
| 2 | North East, PA | 42.22°N | 79.83°W | Jun 2017 | 4 | |
| 3 | Fayetteville, NY | 43.03°N | 76.0°W | Jun 2017 | 4 | |
| 4 | Ashland, OH | 40.87°N | 82.32°W | Jun 2017 | 2 | |
| 7 | Ashland, OH | 40.87°N | 82.32°W | Jul 2016 | 4 | |
| 9 | Amboy, IL | 41.71°N | 89.71°W | Jul 2014 | 4 | 2 |
| 11 | Watertown, NY | 43.97°N | 75.91°W | Jul 2012 | 1 | |
| 12 | Lexington, KY | 38.04°N | 84.50°W | Aug 2012 | 4 | 1 |
| 14 | Canberra, Australia | 35.28°S | 149.13°E | Nov 2007 | 2 | |
| 16 | Hattah, Australia | 34.77°S | 142.39°E | Nov 2005 | 1 | |
| 17 | Curico, Chile | 34.98°S | 71.25°W | 2004 | 2 | 1 |
| 18 | Mildura, Australia | 34.21°S | 142.14°E | Nov 2007 | 1 | |
| 27 | Ithaca, NY | 42.44°N | 76.50°W | Aug 2010 | 4 | |
| 28 | Brooktondale, NY | 42.38°N | 76.39°W | Aug 2017 | 1 | |
| 32 | LeRoy, IL | 40.35°N | 88.76°W | Jul 2017 | 4 | 3 |
| 42 | Watertown, NY | 43.97°N | 75.91°W | Jul 2012 | 4 | |
| 46 | Manhattan, KS | 39.18°N | 96.57°W | May 2015 | 1 | 1 |
| 49 | Břízsko, Czech Republic | 49.54°N | 13.31°E | Jun 2019 | 4 | |

**Table S3.** Sampling localities, GPS Coordinates, and collection information for *P. quatuordecimpunctata* individuals.

| **Locality Number** | **Location** | **Lat** | **Long** | **Date Collected** | **# of Samples Sequenced** | **# of Samples Dropped** |
|---|---|---|---|---|---|---|

| | | | | | | |
|---|---|---|---|---|---|---|
| P2 | Lindenhurst, IL | 42.41°N | 88.03°W | Jun 2017 | 4 | |
| P4 | Watertown, NY | 43.97°N | 75.91°W | Jul 2015 | 2 | |
| P8 | Watertown, NY | 43.97°N | 75.91°W | Jul 2012 | 3 | 1 |
| P9 | Clinton County, IN | 40.28°N | 86.81°W | Aug 2017 | 1 | |
| P10 | LeRoy, IL | 40.35°N | 88.76°W | Jul 2017 | 2 | |
| P19 | Fayetteville, NY | 43.03°N | 76.0°W | Jul 2015 | 2 | |
| P20 | Malone, NY | 44.85°N | 74.29°W | Jul 2016 | 2 | 1 |
| P21 | Ashland, OH | 40.87°N | 82.32°W | Jul 2016 | 2 | 1 |
| P22 | Brooktondale, NY | 42.38°N | 76.39°W | Aug 2017 | 3 | 3 |
| P23 | Watertown, NY | 43.97°N | 75.91°W | Aug 2015 | 4 | 4 |
| P25 | Muncie, IN | 40.19°N | 85.39°W | May 2012 | 3 | 3 |
| P26 | Ashland, OH | 40.87°N | 82.32°W | Jul 2011 | 1 | 1 |
| P27 | Amboy, IL | 41.71°N | 89.33°W | Aug 2014 | 3 | 3 |
| P28 | Fayetteville, NY | 43.03°N | 76.0°W | Jun 2019 | 3 | 3 |
| P29 | Watertown, NY | 43.97°N | 75.91°W | Jun 2019 | 3 | 1 |
| P30 | Preble, NY | 42.74°N | 76.15°W | Jun 2019 | 3 | |
| P31 | State College, PA | 40.79°N | 77.86°W | Jun 2010 | 4 | |
| P32 | Břízsko, Czech Republic | 49.54°N | 13.31°E | Jun 2019 | 4 | |

**Table S4.** Pairwise $F_{ST}$ values as calculated by StAMPP in 16 populations of *H. variegata*. Values in bold are significant at the 0.05 level. Pairwise $F_{ST}$ was not able to be calculated between two populations that contain a single individual and are notated as "*NA*".

| | ***11*** | ***12*** | ***14*** | ***16*** | ***17*** | ***18*** | ***2*** | ***27*** | ***28*** | ***3*** | ***32*** | ***4*** | ***42*** | ***49*** | ***7*** | ***9*** |
|---|---|---|---|---|---|---|---|---|---|---|---|---|---|---|---|---|
| ***11*** | -- | 0.032 | **0.29** | *NA* | *NA* | *NA* | -0.002 | 0.015 | *NA* | 0.008 | 0.036 | -0.045 | -0.01 | **0.131** | -0.025 | -0.101 |
| ***12*** | 0.032 | -- | **0.182** | **0.082** | **0.076** | **0.1** | -0.005 | 0.019 | 0.039 | **0.029** | **0.034** | 0.03 | 0.013 | **0.067** | 0.025 | 0.01 |
| ***14*** | **0.29** | **0.182** | -- | -0.022 | **0.157** | -0.018 | **0.14** | **0.149** | **0.243** | **0.156** | **0.153** | **0.176** | **0.144** | **0.179** | **0.191** | **0.18** |
| ***16*** | *NA* | **0.082** | -0.022 | -- | *NA* | *NA* | 0.039 | **0.062** | *NA* | **0.059** | **0.076** | 0.017 | **0.076** | **0.078** | **0.09** | -0.045 |
| ***17*** | *NA* | **0.076** | **0.157** | *NA* | -- | *NA* | 0.047 | **0.077** | *NA* | **0.096** | **0.095** | 0.046 | **0.076** | **0.134** | 0.036 | 0.036 |
| ***18*** | *NA* | **0.1** | -0.018 | *NA* | *NA* | -- | **0.09** | **0.104** | *NA* | **0.095** | **0.127** | 0.015 | **0.111** | **0.149** | **0.114** | 0.013 |
| ***2*** | -0.002 | -0.005 | **0.14** | 0.039 | 0.047 | **0.09** | -- | -0.004 | 0.001 | -0.018 | -0.002 | 0.022 | -0.003 | **0.033** | 0 | 0.008 |
| ***27*** | 0.015 | 0.019 | **0.149** | **0.062** | **0.077** | **0.104** | -0.004 | -- | -0.002 | 0.008 | -0.011 | 0 | -0.01 | **0.031** | -0.022 | 0.026 |
| ***28*** | *NA* | 0.039 | **0.243** | *NA* | *NA* | *NA* | 0.001 | -0.002 | -- | 0.036 | 0.027 | -0.033 | 0.01 | **0.073** | -0.017 | 0.01 |
| ***3*** | 0.008 | **0.029** | **0.156** | **0.059** | **0.096** | **0.095** | -0.018 | 0.008 | 0.036 | -- | -0.032 | 0.016 | 0.008 | **0.052** | 0.013 | 0.026 |
| ***32*** | 0.036 | **0.034** | **0.153** | **0.076** | **0.095** | **0.127** | -0.002 | -0.011 | 0.027 | -0.032 | -- | 0.019 | 0 | **0.039** | 0.006 | 0.029 |
| ***4*** | -0.045 | 0.03 | **0.176** | 0.017 | 0.046 | 0.015 | 0.022 | 0 | -0.033 | 0.016 | 0.019 | -- | 0.002 | **0.054** | -0.02 | -0.03 |
| ***42*** | -0.01 | 0.013 | **0.144** | **0.076** | **0.076** | **0.111** | -0.003 | -0.01 | 0.01 | 0.008 | 0 | 0.002 | -- | **0.05** | -0.012 | 0.013 |
| ***49*** | **0.131** | **0.067** | **0.179** | **0.078** | **0.134** | **0.149** | **0.033** | **0.031** | **0.073** | **0.052** | **0.039** | **0.054** | **0.05** | -- | **0.05** | **0.072** |
| ***7*** | -0.025 | 0.025 | **0.191** | **0.09** | 0.036 | **0.114** | 0 | -0.022 | -0.017 | 0.013 | 0.006 | -0.02 | -0.012 | **0.05** | -- | 0.008 |
| ***9*** | -0.101 | 0.01 | **0.18** | -0.045 | 0.036 | 0.013 | 0.008 | 0.026 | 0.01 | 0.026 | 0.029 | -0.03 | 0.013 | **0.072** | 0.008 | -- |

**Table S5**. Pairwise $F_{ST}$ values as calculated by StAMPP in 12 populations of *P. quatuordecimpunctata*. Values in bold are significant at the 0.05 level. Pairwise $F_{ST}$ was not able to be calculated between two populations that contain a single individual and are notated as "*NA*".

| | ***P10*** | ***P19*** | ***P2*** | ***P20*** | ***P21*** | ***P29*** | ***P30*** | ***P31*** | ***P32*** | ***P4*** | ***P8*** | ***P9*** |
|---|---|---|---|---|---|---|---|---|---|---|---|---|
| ***P10*** | - | **0.070** | **0.034** | 0.023 | **0.072** | 0.035 | 0.025 | **0.036** | **0.164** | 0.021 | **0.040** | 0.050 |
| ***P19*** | **0.070** | - | **0.021** | -0.009 | 0.032 | **0.037** | 0.021 | 0.012 | **0.121** | 0.007 | **0.032** | 0.013 |
| ***P2*** | **0.034** | **0.021** | - | 0.008 | 0.026 | **0.033** | 0.012 | **0.015** | **0.074** | 0.008 | -0.008 | -0.004 |
| ***P20*** | 0.023 | -0.009 | 0.008 | - | -0.004 | 0.010 | -0.012 | 0.019 | **0.13** | -0.033 | 0.010 | -0.06 |
| ***P21*** | **0.072** | 0.032 | 0.026 | -0.004 | - | **0.075** | 0.026 | 0.03 | **0.162** | -0.010 | **0.074** | *NA* |
| ***P29*** | 0.035 | **0.037** | **0.033** | 0.01 | **0.075** | - | 0.037 | 0.033 | **0.164** | 0.019 | **0.039** | **0.047** |
| ***P30*** | 0.025 | 0.021 | 0.012 | -0.012 | 0.026 | **0.037** | - | 0.004 | **0.1** | 0.019 | 0.001 | -0.023 |
| ***P31*** | **0.036** | 0.012 | 0.015 | 0.019 | 0.030 | **0.033** | 0.004 | - | **0.063** | **0.030** | -0.001 | 0.008 |
| ***P32*** | **0.164** | **0.121** | **0.074** | **0.130** | **0.162** | **0.164** | **0.100** | **0.063** | - | **0.120** | **0.124** | **0.150** |
| ***P4*** | 0.021 | 0.007 | 0.008 | -0.033 | -0.010 | 0.019 | 0.019 | 0.030 | **0.120** | - | -0.005 | -0.022 |
| ***P8*** | **0.040** | **0.032** | -0.008 | 0.010 | **0.074** | **0.039** | 0.001 | -0.001 | **0.124** | -0.005 | - | 0.031 |
| ***P9*** | 0.050 | 0.013 | -0.004 | -0.060 | *NA* | **0.047** | -0.023 | 0.008 | **0.150** | -0.022 | 0.031 | - |

**Table S6.** Estimates of contemporary migration rates, 95% confidence intervals from BayesAss3-SNPs using the *P. quatuordecimpunctata* ddRADseq data. Population 0 refers to the USA samples, while 1 is the Czech Republic. m[0][1] would thus represent the estimated migration rate (here the proportion of individuals with migrant ancestry in every generation) from the USA to the Czech Republic. All estimates had an ESS of >10,497, out of a total of 10,596 samples.

| **Parameter** | **Estimate (mean)** | **95% HPD Interval** |
|---|---|---|
| m[0][0] | 0.9877 | [0.8977, 1] |
| m[0][1] | 0.0123 | [4.5e-7, 0.036] |
| m[1][0] | 0.0565 | [1.47e-5, 0.1525] |
| m[1][1] | 0.9435 | [0.8475, 1] |

**Table S7.** Estimates of contemporary migration rates, 95% confidence intervals from BayesAss3-SNPs using the Hvar ddRADseq data. Population 0 refers to the USA samples, while 1 includes samples from Australia and Chile, and 2 for samples from the Czech Republic. m[0][1] would thus represent the estimated migration rate (here the proportion of individuals with migrant ancestry in every generation) from the USA to the Australia + Chile cluster. All estimates had an ESS of >43,878, out of a total of 44,470 samples.

| **Parameter** | **Estimate (mean)** | **95% HPD Interval** |
|---|---|---|
| m[0][0] | 0.917 | [0.8242, 0.9953] |
| m[0][1] | 0.042 | [3.41e-7, 0.115] |
| m[0][2] | 0.042 | [6.82e-7, 0.116] |
| m[1][0] | 9.53e-3 | [8.83e-8, 0.0279] |
| m[1][1] | 0.981 | [0.955, 0.999] |
| m[1][2] | 9.47e-4 | [2.13e-8, 0.0278] |
| m[2][0] | 0.0473 | [7.97e-7, 0.1292] |
| m[2][1] | 0.0474 | [6.43e-7, 0.1307] |
| m[2][2] | 0.9053 | [0.8035, 0.9933] |

**Table S8**. Egg, larval, pupal and pre-imaginal (egg to adult) developmental times (days) (Mean ± SD) for *Propylea quatuordecimpunctata* and *Hippodamia variegata* at four constant photoperiod conditions (L:D 16:8. 14:10, 12:12, 10:14) at 22°C. Populations collected in Jefferson County, NY. F1 laboratory reared individuals on pea aphids (*Acyrthosiphon pisum*) and *Ephestia kuehniella* eggs. One-way ANOVA for effect of photoperiod on developmental times. *Hippodamia variegata* data from Obrycki (2018).

| **Population Developmental Period** | **Daylength Treatment (L:D)** | | | | **ONE-WAY ANOVA**<br>df = 3,157 |
|---|---|---|---|---|---|
| | **16:8** | **14:10** | **12:12** | **10:14** | |
| *Propylea quatuordecimpunctata*<br>Egg<br>Larval<br>Pupal<br>Pre-imaginal | N = 38<br><br>4.3 ± 0.5<br>11.8 ± 2.0<br>4.9 ± 0.4<br>21.1 ± 2.3 | N = 45<br><br>3.2 ± 0.4<br>11.0 ± 0.6<br>5.0 ± 0.3<br>19.2 ± 0.6 | N = 40<br><br>3.9 ± 0.4<br>10.2 ± 0.4<br>5.0 ± 0.3<br>19.0 ± 0.7 | N = 38<br><br>4.0 ± 0.0<br>10.4 ± 0.9<br>4.1 ± 0.7<br>18.6 ± 0.6 | <br><br>F = 66.23, P < 0.0001<br>F =15.85, P < 0.0001<br>F = 31.62, P < 0.0001<br>F = 30.72, P < 0.0001 |
| *Hippodamia variegata*<br>Egg<br>Larval<br>Pupal<br>Pre-imaginal | N = 26<br><br>3.2 ± 0.4<br>13.9 ± 1.4<br>5.1 ± 0.5<br>22.2 ± 1.4 | N = 34<br><br>3.9 ± 0.4<br>12.9 ± 0.6<br>4.9 ± 0.3<br>21.6 ± 0.6 | N = 29<br><br>4.1 ± 0.8<br>11.7 ± 1.0<br>4.4 ± 0.5<br>20.2 ±1.6 | N =36<br><br>3.4 ± 0.5<br>12.3 ± 0.9<br>4.5 ± 0.5<br>20.2 ± 1.3 | ONE-WAY ANOVA<br>df = 3, 121<br>F = 17.03, P < 0.0001<br>F = 25.21, P < 0.0001<br>F = 15.22, P < 0.0001<br>F = 19.78, P < 0.0001 |

**Table S9**. Two-way ANOVA table for egg, larval, pupal, and total preimaginal development of *Propylea quatuordecimpunctata* and *Hippodamia variegata* at constant photoperiods: L:D 16:8, 14:10, 12:12, 10:14 at 22° ±1°C. Species collected in Jefferson County, New York.

| **Developmental Period** | **Source** | **df** | **Sum of Squares** | **F ratio** | **Prob > F** |
|---|---|---|---|---|---|
| Egg | Model | 7 | 43.207 | 30.55 | < 0.0001 |
| | Species | 1 | 2.444 | 12.10 | 0.0006 |
| | Daylength | 3 | 7.650 | 12.62 | < 0.0001 |
| | Species x Daylength | 3 | 31.997 | 52.79 | < 0.0001 |
| Larval | Model | 7 | 352.273 | 44.05 | < 0.0001 |
| | Species | 1 | 226.890 | 198.58 | < 0.0001 |
| | Daylength | 3 | 133.265 | 38.88 | < 0.0001 |
| | Species x Daylength | 3 | 2.795 | 0.82 | 0.4862 |
| Pupal | Model | 7 | 30.135 | 20.03 | < 0.0001 |
| | Species | 1 | 0.0035 | 0.02 | 0.8986 |
| | Daylength | 3 | 22.758 | 35.30 | < 0.0001 |
| | Species x Daylength | 3 | 7.417 | 11.51 | < 0.0001 |
| Preimaginal (egg to adult) | Model | 7 | 406.505 | 37.49 | < 0.0001 |
| | Species | 1 | 179.174 | 115.67 | < 0.0001 |
| | Daylength | 3 | 207.568 | 44.67 | < 0.0001 |
| | Species x Daylength | 3 | 17.50 | 3.77 | 0.0112 |

**Table S10**. Pre-oviposition period (days) as a measure of induction and duration of adult diapause in *Propylea quatuordecimpunctata* (P-14) and *Hippodamia variegata*. Constant photoperiods: L:D 16:8; 14:10; 12:12; and 10:14 at 22°C ±1°C. Species collected in Jefferson County, NY. Pre-oviposition period: Mean (days ± SE); Min-Max days; Median (days); [N] = number of ovipositing females. Non-parametric log-rank analysis: (1) within row examines response to the 4 photoperiods. *Hippodamia variegata* data from (Obrycki 2018).

| **Population** | **Daylength Treatment (L:D)** | | | | |
|---|---|---|---|---|---|
| | **16:8** | **14:10** | **12:12** | **10:14** | **Log-Rank**<br>ChiSq,df, P |
| P-14<br>Mean ± SE<br>Min-Max [N]<br>Median | <br>12.1 ± 4.8<br>7-24 [13]<br>10 | <br>-<br>- [0]<br>- | <br>-<br>- [0]<br>- | <br>-<br>- [0]<br>- | 89.21,<br>df=3,<br>P<. 0.0001 |
| *H. variegata*<br><br>Mean ± SE<br>Min-Max [N]<br>Median | <br><br>11.3 ± 2.1<br>5-26 [13]<br>7.5 | <br><br>56.2 ± 4.6<br>25-69 [8]<br>32 | <br><br>61.3 ± 0.5<br>58-62 [3]<br>58 | <br><br>68.7 ± 0.3<br>65-69 [3]<br>68 | 77.46,<br><br>df =3,<br><br>P< 0.0001 |
| **Log-Rank**<br>ChiSq,df, P | 0.076, df = 3,<br>P = 0.7824 | 10.85, df = 3,<br>P = 0.0010 | 4.36, df=3,<br>P = 0.0367 | 2.79, df=3,<br>P = 0.095 | |

**Table S11**. Pre-oviposition period (days) as a measure of induction and duration of adult diapause in *Propylea quatuordecimpunctata* (P-14) from Jefferson County, New York, USA and *Propylea quatuordecimpunctata* from Montreal, Quebec, Canada (P-14 CAN). Constant photoperiods: L:D 16:8; 14:10; 12:12; and 10:14 at 22°C ±1°C. Pre-oviposition period: Mean (days ± SE); Min-Max days; Median (days); [N] = number of ovipositing females. Non-parametric log-rank analysis: (1) within row examines response to the 4 photoperiods (2) within columns compares responses of females from the two populations to the same photoperiod. *Propylea quatuordecimpunctata* data from Montreal, Quebec, Canada from (Obrycki et al 1993).

| **Population** | **Daylength Treatment (L:D)** | | | | |
|---|---|---|---|---|---|
| | **16:8** | **14:10** | **12:12** | **10:14** | **Log-Rank** ChiSq,df, P |
| P-14<br>Mean ± SE<br>Min-Max [N]<br>Median | <br>12.1 ± 4.8<br>7-24 [13]<br>10 | <br>-<br>- [0]<br>- | <br>-<br>- [0]<br>- | <br>-<br>- [0]<br>- | 89.21,<br>df=3,<br>P<. 0.0001 |
| P-14 Canada<br><br>Mean ± SE<br>Min-Max [N]<br>Median | <br><br>17.2 ± 21.4<br>7-73 [9]<br>9 | <br><br>8.5 ± 0.7<br>8-9 [2]<br>8.5 | <br><br>9.0 ± 0.0<br>9 [1]<br>- | <br><br>8.0 ± 0.0<br>8 [1]<br>- | 25.73, df =3,<br>P< 0.0001 |
| **Log-Rank**<br>ChiSq,df, P | 0.33, df = 1,<br>P = 0.5634 | 3.58, df = 1,<br>P = 0.0583 | 1.60, df=1,<br>P = 0.2059 | 1.30, df=1,<br>P = 0.2542 | |

**Table S12**. Percentage of female *Propylea quatuordecimpunctata* (P-14) and *Hippodamia variegata* from Jefferson County, NY and *P. quatuordecimpunctata* (P-14 CAN) females from Montreal, Quebec, Canada in diapause at four constant photoperiods: L:D 16:8; 14:10; 12:12; and 10:14 at 22°C ±1°C. The criteria for reproductive diapause in a female was 2X the median pre-oviposition period (days) observed at L:D 16:8 for each species / population.

| **Species / Population** | **Daylength Treatment (L:D)** | | | | **Reference** |
|---|---|---|---|---|---|
| | **16:8** | **14:10** | **12:12** | **10:14** | |
| P-14<br>USA, New York<br>Jefferson Cty | 8 %<br>2X Median = 20 days | 100 % | 100 % | 100 % | Present Study |
| P-14 CAN<br>Canada, Quebec<br>Montreal | 22 %<br>2X Median = 18 days | 80 % | 90 % | 90 % | Obrycki et al. 1993 |
| *H. variegata*<br>USA, New York<br>Jefferson Cty | 15 %<br>2X Median = 15 days | 100 % | 100 % | 100 % | Obrycki 2018 |

**Figure S1.** Full sampling map of **(A)** *Hippodamia variegata* and **(B)** *Propylea quatuordecimpunctata.*

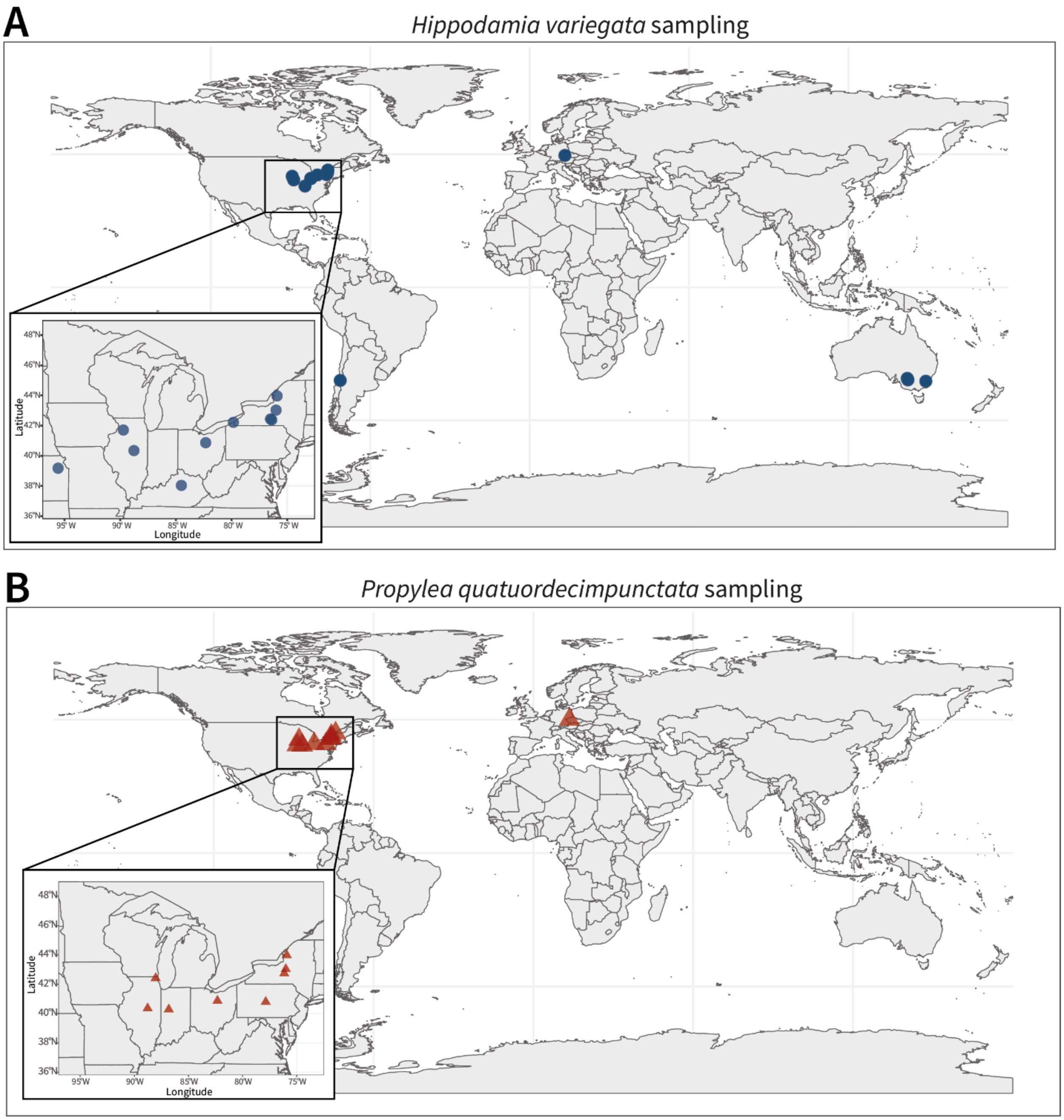

**Figure S2.** Additional summary plots from DAPC and ADMIXTURE analyses. DAPC cluster assignment plots (**A**) with DAPC Bayesian Information Criterion (BIC) and ADMIXTURE cross-validation (CV) comparison for all *H. variegata* individuals (**B**) and U.S. *H. variegata* individuals (**C**). DAPC cluster assignment plots (**D**) with DAPC Bayesian Information Criterion (BIC) and ADMIXTURE cross-validation (CV) comparison for all *P. quatuordecimpunctata* individuals (**E**) and U.S. *P. quatuordecimpunctata* individuals (**F**).

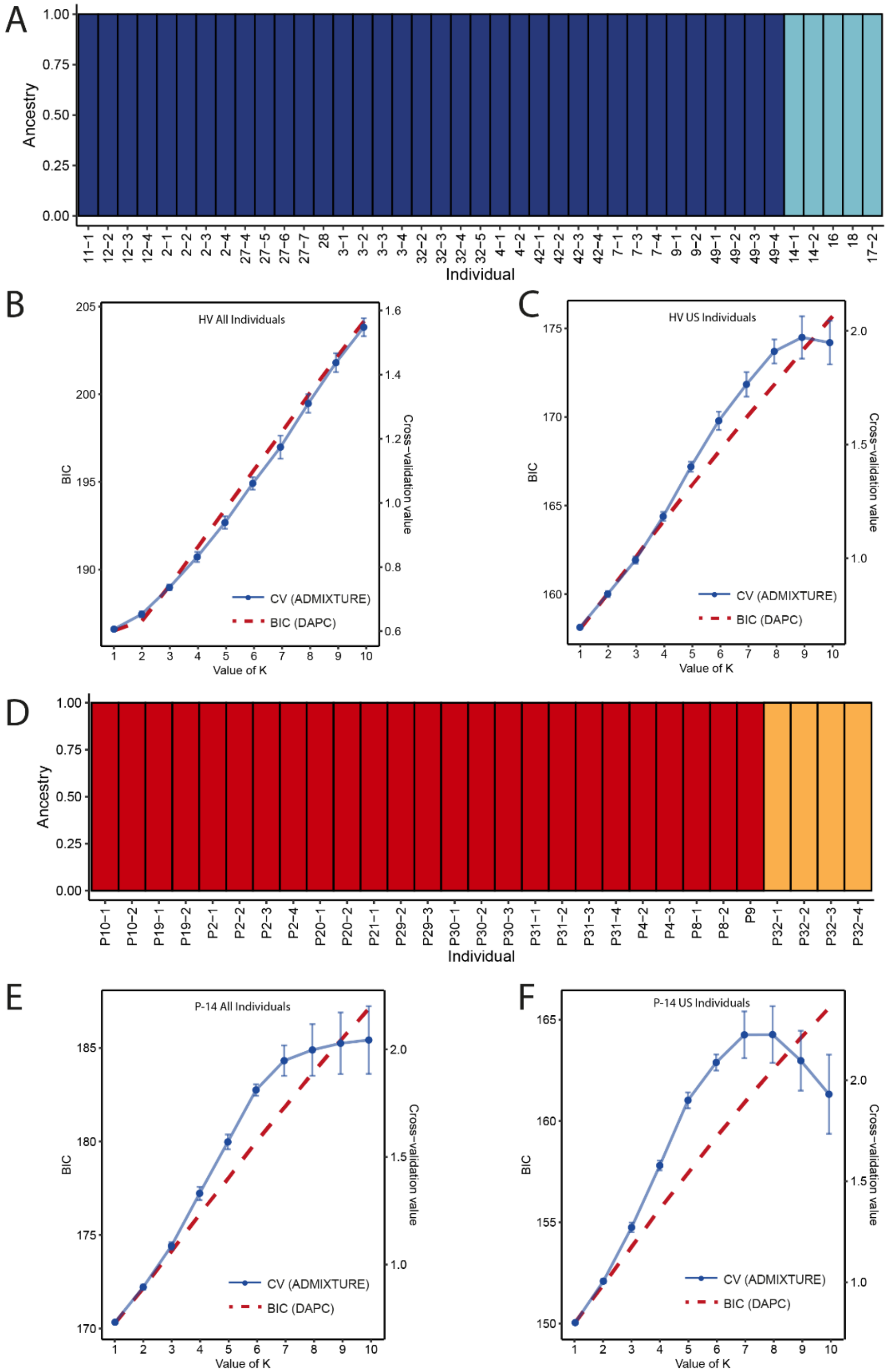

**Figure S3.** Additional summary plots from conStruct analyses. Layer contribution plots from conStruct runs considering allele frequencies of individual samples of (**A**) *H. variegata* (HV) and (**B**) *P. quatuourdecimpunctata* (P14). Comparison of predictive accuracy of nonspatial and spatial conStruct models across different *K*s in construct runs considering average allele frequencies in populations of (**C**) HV and (**D**) P14. Layer contribution plots from conStruct runs considering average allele frequencies in populations of (**E**) HV and (**F**) P14.

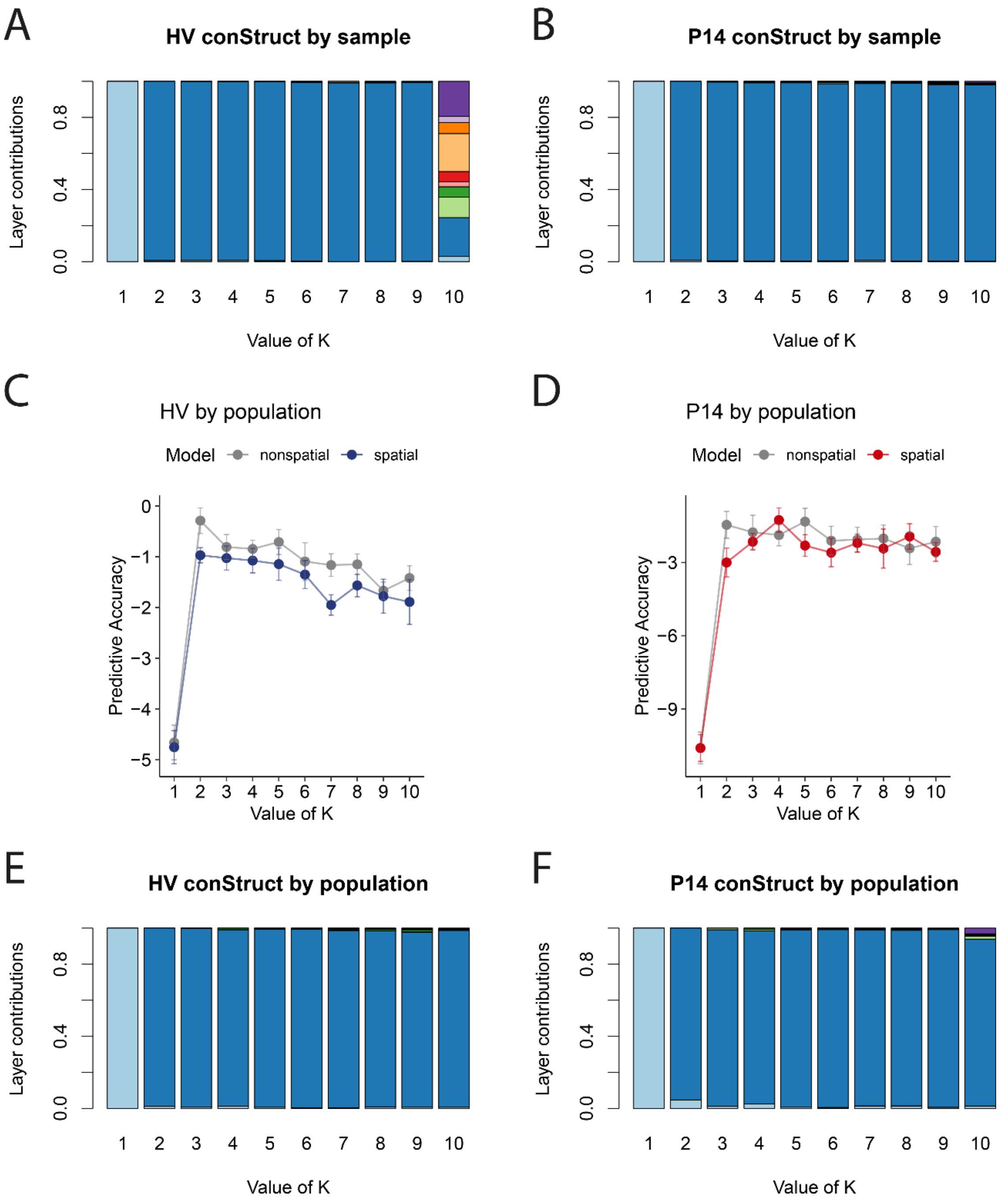

**Figure S4.** Global and local structure eigenvalues from sPCA analyses of (**A**) *H. variegata* (HV) and (**B**) *P. quatuourdecimpunctata* (P14).

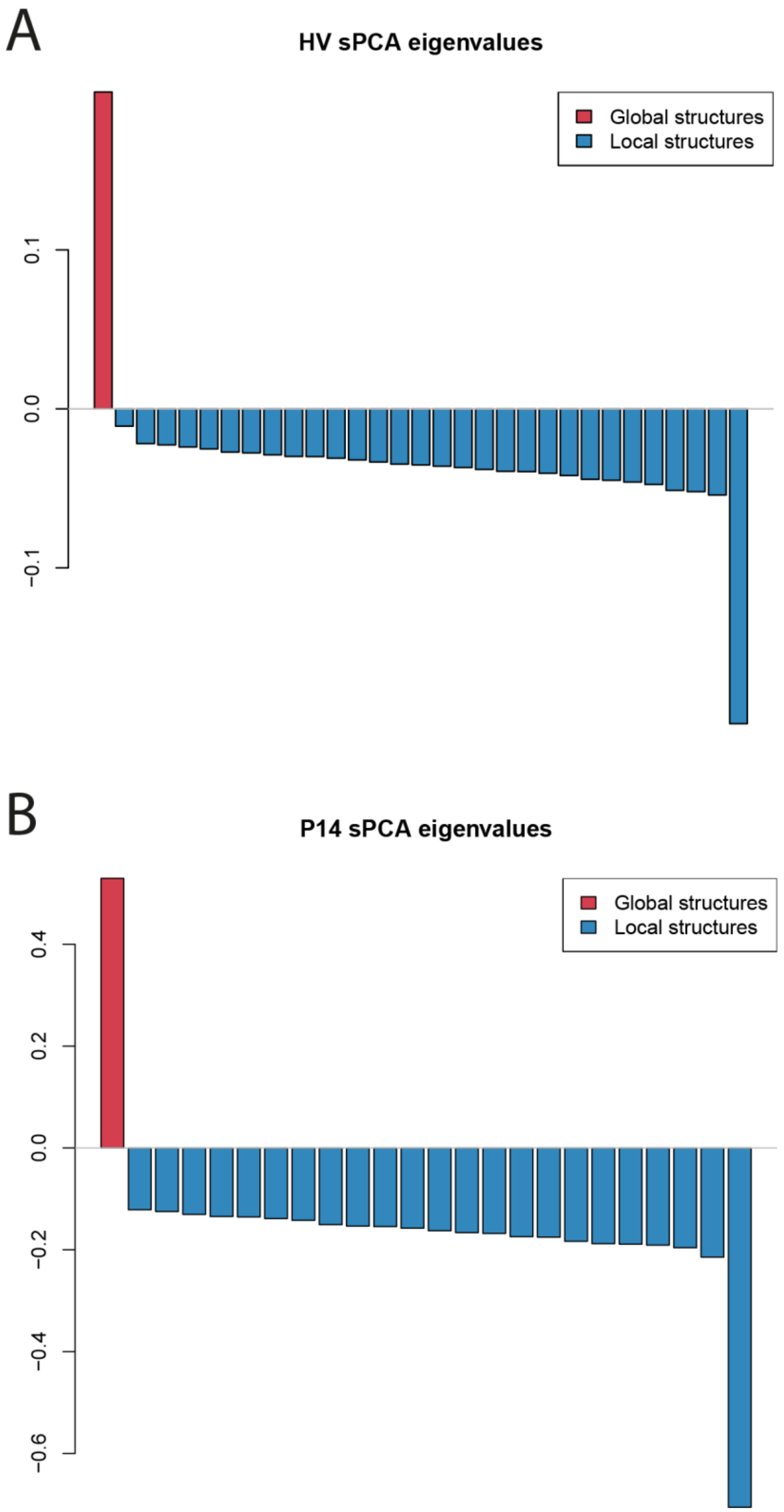
